# Sweet to arrive, fermented to fly: flower-insect interactions in a Mexican dry forest population of *Annona macroprophyllata* (Annonaceae)

**DOI:** 10.1101/2025.07.17.665288

**Authors:** Yuridia Esmeralda Llaven-Albores, Iván de-la-Cruz-Chacón, Alma Rosa Gonzalez-Esquinca, Oscar Pérez Flores, Marisol Castro-Moreno, Andres Ernesto Ortiz-Rodriguez

## Abstract

Flower scents are crucial for attracting pollinators to Annonaceae flowers. Pollinators are likely drawn exclusively to female flowers. Since pollinators remain inside the pollination chamber throughout anthesis in several species, it is believed that the aroma during the male phase does not attract pollinators and may even repel them. However, few studies have characterised the scents of Annonaceae species’ flowers during both the male and female phases, so this hypothesis has not been comprehensively evaluated. Here we characterise the reproductive ecology of *Annona macroprophyllata*, including individual- and flower-level phenology, mutualistic and non-pollinating interactions, and floral scent emissions. Our results showed that *Annona macroprophyllata* has receptive flowers at night, with the entire extent of reproductive activity restricted to an 8 hours-period. Female and male flowers are synchronous with differences in the proportion of flowers depending on the nighttime. Low diversity and abundance of floral visitors were observed. Four guilds were distinguished as pollinators, florivores, insect predators and random or non-specific visitors. Pollinators were insects (Nitulidae) that entered the pollination chamber during the female phase and remained inside the flower throughout anthesis. The comparison of floral scent diversity among flower phases showed that scent composition differs between the phases of *A. macroprophyllata* flowers. The scent of the female flower mimics the sweet smell of fruit, which usually attracts Nitulid beetles. In contrast, the fermented odors of the male phase seem to repel them, contributing to their release or ensuring that they visit only female flowers. This study reveals an attraction-repulsion pollinator system in *Annona macroprophyllata*, which could be a common floral attribute within the Annonaceae family.

## Introduction

The Annonaceae family, a species-rich lineage within the Magnoliales order, comprises approximately 2,500 species of trees, shrubs, and lianas, distributed across 110 genera. This family has a pantropical distribution and is one of the most representative lineages of woody plants in tropical forests. Flowers in Annonaceae are typically trimerous, hermaphrodite and protogynous, since they are self-compatible, their species have developed several mechanisms that promote cross-pollination. More specifically, a sexually non-functional interrim phase temporarily separates the male and female phases of the flower and prevents self-pollination [1–2]. Cross-pollination in Annonaceae is promoted by insects and pollination by flies, cockroaches, thrips, bees, and small and large beetles has been reported [3–4]. Pollinators are attracted to the flowers by scent, thermogenesis or the production of stigmatic secretions [5]. Also, the fleshy inner petals of several species are frequently used as mating sites [3–4]. Recently, it has been shown that the duration and phenology of Annonaceae flowers are coupled to the circadian rhythm of pollinators [6]. Furthermore, the floral structure and its arrangement during the anthesis plays a very important role in the identity and behaviour of Annonaceae pollinators [7].

Among the Annonaceae genera, *Annona* species have a highly specialised pollination strategy. *Annona* flowers do not fully spread their petals during anthesis, forming a flower chamber (= pollination chamber) that serves as a size filter mechanism for visiting beetles, which are attracted to the flowers by their strong scents and the presence of food rewards like pollen and floral tissues [7–9]. The pollinators (insects that manage to pass through the pollination chamber filter) enter the flower during the female phase and remain there for the rest of the anthesis [10]. Then all the insects inside the pollination chamber, often carrying pollen on their bodies, are suddenly released from the flower by dropping the petals [10]. In this pollination strategy, flower scents play a crucial role in attracting pollinators to *Annona* flowers [5,11]. The emission of these scents signals the receptivity of the female phase. Thus, most studies on the genus *Annona* report sweet, fruity odors, or less frequently, unpleasant odors in the flowers during anthesis [11]. Experimental studies indicate that *Annona* pollinators are attracted exclusively to female flowers.

Given that pollinators remain inside the pollination chamber throughout anthesis, it is assumed that the aroma during the male phase does not attract pollinators to the flowers [12]. However, to our knowledge, there are no studies characterising the odors of *Annona* flowers in the male and female phases. If differences are found, it would support the hypothesis that only female flowers produce odors that attract pollinators. This odor changes towards the end of anthesis, promoting beetles with pollen loads to visit only female flowers of other individuals [12].

Here we characterise the reproductive ecology of *Annona macroprophyllata* Donn Sm., including individual- and flower-level phenology, mutualistic and non-pollinating interactions, and floral scent emissions. *Annona macroprophyllata, c*ommonly known as “Papausa” or “Ilama”, is an edible fruit that naturally grows in the dry forests of Mexico and Central America (Fig. 1). Due to its commercial and edible value, it is often cultivated in backyards and used as part of living fences. While some information about its vegetative and reproductive phenology is available [13], its interactions with floral visitors have not yet been studied. Our specific objectives were to: (1) identify the start and end of the male and female phases of the flower, (2) assess the diversity of floral visitors during each reproductive phase, and (3) characterise the chemical compounds responsible for the flower’s aromas in both reproductive phases.

**Fig. 1.**
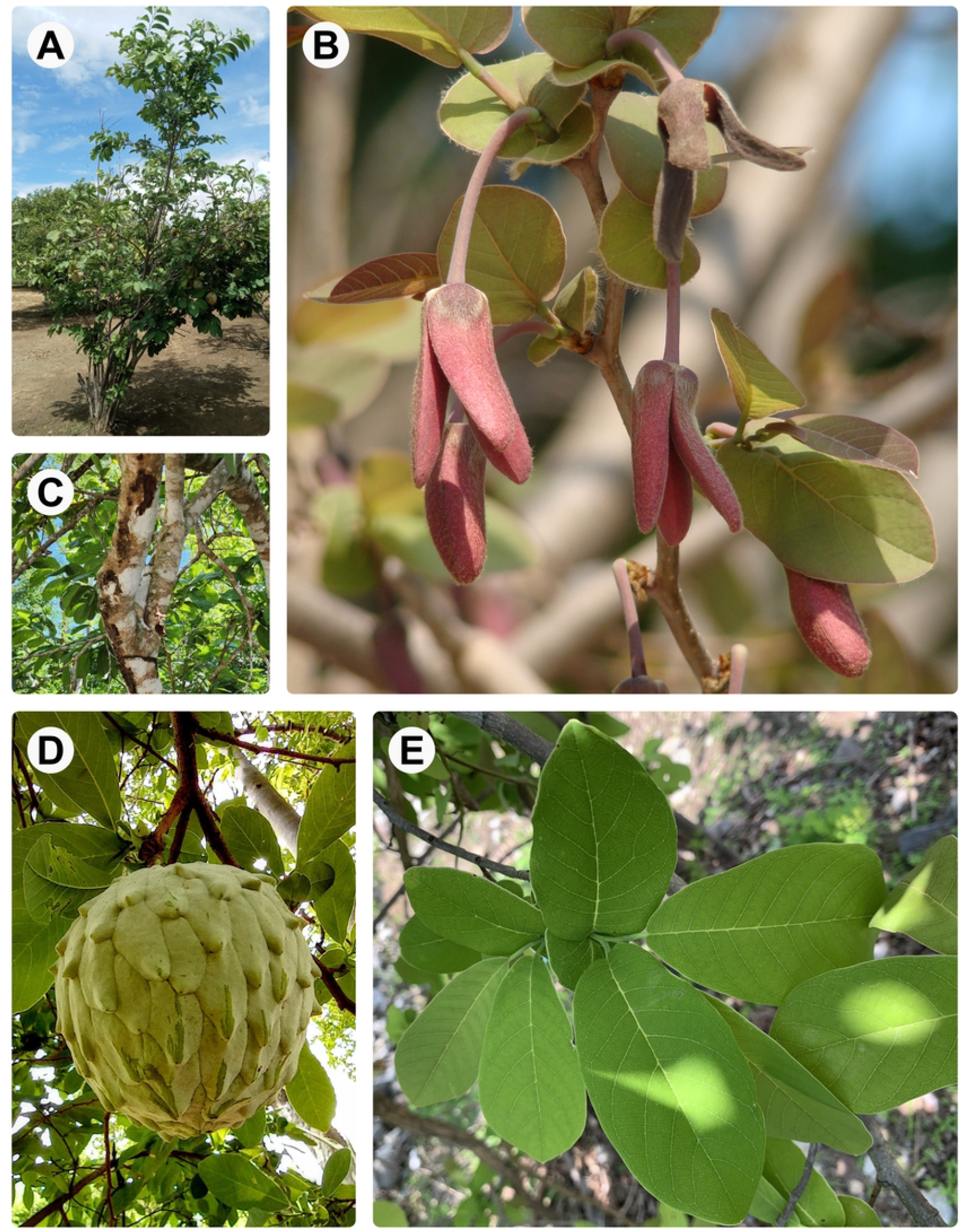
*Annona macroprophyllata* (Annonaceae). A) habit, B) reproductive twigs with flower buds and mature flowers, C) main trunk and bark, D) Fruit, E) Leaves. Photographs by A) Cándido Hernández Aguilar, B) Ivan De la Cruz, C) Carlos G. Velazco-Macias, D) José Rosario Soto, E) alana_atl, available at inaturalist.org.

## Materials and methods

### Study site

The study site is located in Chiapas, Mexico, between the municipalities of Chiapa de Corzo and Parral, at coordinates 16°31’49.0” N and 92°58’06.7” W. This agroecosystem features patches of natural dry forest interspersed with cultivated tree species such as Jocote (*Spondias purpurea* L.), Nance (*Byrsonima crassifolia* (L.) Kunth), and Chincuya (*Annona purpurea* Moc. & Sessé ex Dunal).

### Floral phenology and floral visitors’ diversity

Ten reproductive *Annona macroprophyllata* trees were selected, with distances between them ranging from four to 40 meters. Full flowering was observed from April to May in 2017. The trees did not exhibit synchronous flowering, so weekly field trips were scheduled to monitor the flowering of the selected trees. Specifically, the abundance of mature flowers per week in each tree was determined. The morphology of the mature flowers, length of the petals and diameter of the flower base, were measured on three different trees (10 flowers per tree).

Flower-level phenology was determined on the four different tree individuals (40 flowers observed in total). We carried out two 24-hour observation periods, to determine the timing of the onset of stigmatic receptivity (female phase), its duration and the start and duration of the staminal phase (male phase). The presence or absence of stigmatic fluid in female and male flowers and the receptivity of the carpels were evaluated using the peroxidase enzyme reaction method [14]. The identity of the visitor, visitor behavior and stage of sexual maturity of the flower were recorded. External visitors and visitors inside the flower were captured when possible. Pollinating insects were identified as those that entered the pollination chamber and remained there throughout anthesis.

### Flowers scents characterization

Flower scent characterization was carried out at Universidad de Ciencias y Artes de Chiapas, Mexico. Flowers in pistillate (female) and staminate (male) phases were collected in new polypropylene bags (15 × 15 cm). The bags were sealed to limit air movement. Steam distillation using Clevenger-type machine was immediately performed for six hours. The essential oil was collected and dissolved in hexane for analysis, which was done in triplicate. The samples were analysed using gas chromatography-mass spectrometry (GC-MS) on a Perkin Elmer Clarus No.680-5Q8T system, equipped with an elite 1 column. The samples dissolved in 2 mL of hexane were injected with a split ratio of 50. The carrier gas was helium at 1 mL/min. The injector temperature was 250°C, the transfer line was 250°C, and the detector was set to 280°C. The elution time was 30 minutes per sample. The NIST 98 data library (NIST algorithm) was used for component identification [15].

## Results

### Floral phenology and floral visitors’ diversity

During the first week of observation (April 30 - May 5), only four trees had mature flowers, a total of 169 flowers, 42 flowers on average per tree. During the second week (May 6 - May 12), the highest flowering peak was observed at the study site, seven of the ten studied trees had mature flowers, a total of 3035 flowers, 433 flowers on average per tree. During the last week of observation (May 13 - May 20), all the trees had mature flowers, a total of 193 mature flowers, still the number of available flowers decreased considerably, with an average of 19 mature flowers per tree. Flower size is homogeneous among individuals, with an average petal length of 3.18 cm ± 0.37 (2.5‒3.9 cm; N= 30), and an average diameter of the flower base of 1.49 cm ± 0.53 (0.6‒3.0 cm; N= 30).

The flowers were markedly protogynous, with the entire extent of reproductive activity restricted to an 8 hours-period (Fig. 2). The reproductive phases (female and male phases) are synchronous with differences in the proportion of flowers in each phase depending on the nighttime (Fig. 2). At the beginning of the afternoon, between 1500 and 1600 hours, 25 % of the flowers observed started the female phase. Thus, the three red petals of the flowers closed so they formed the pollination chamber over the reproductive organs, the stigmas seem enlarged, and were covered by a transparent exudate so they had a wet appearance; this phase was correlated with the emission of a strong sweet-like odour, with positive peroxide test and with a further increase in floral visitors. Afterwards, between 1700 and 1800 hours, 50% of the flowers observed were in the female phase and 25% in the male phase, showing the synchrony between phases (Fig. 2). During male phase the petals are more distant from each other and the pollination chamber is open, the anthers dehisced, carpels look dry and reddened, and there was an obvious change in floral scent. The male phase ends with the detachment of the petals and the fall of all the stamens. Then, between 1900 and 2100 hours, 75% of the flowers observed were in the female phase and 25% in the male phase, showing the highest peak of stigmatic receptivity among the tres (Fig. 2). Lastly, between 2130 and 2400 hours, 75% of the flowers observed were in the male phase and 25% in the female phase, with many senescent flowers, thus showing the end of anthesis (Fig. 2).

**Fig. 2.**
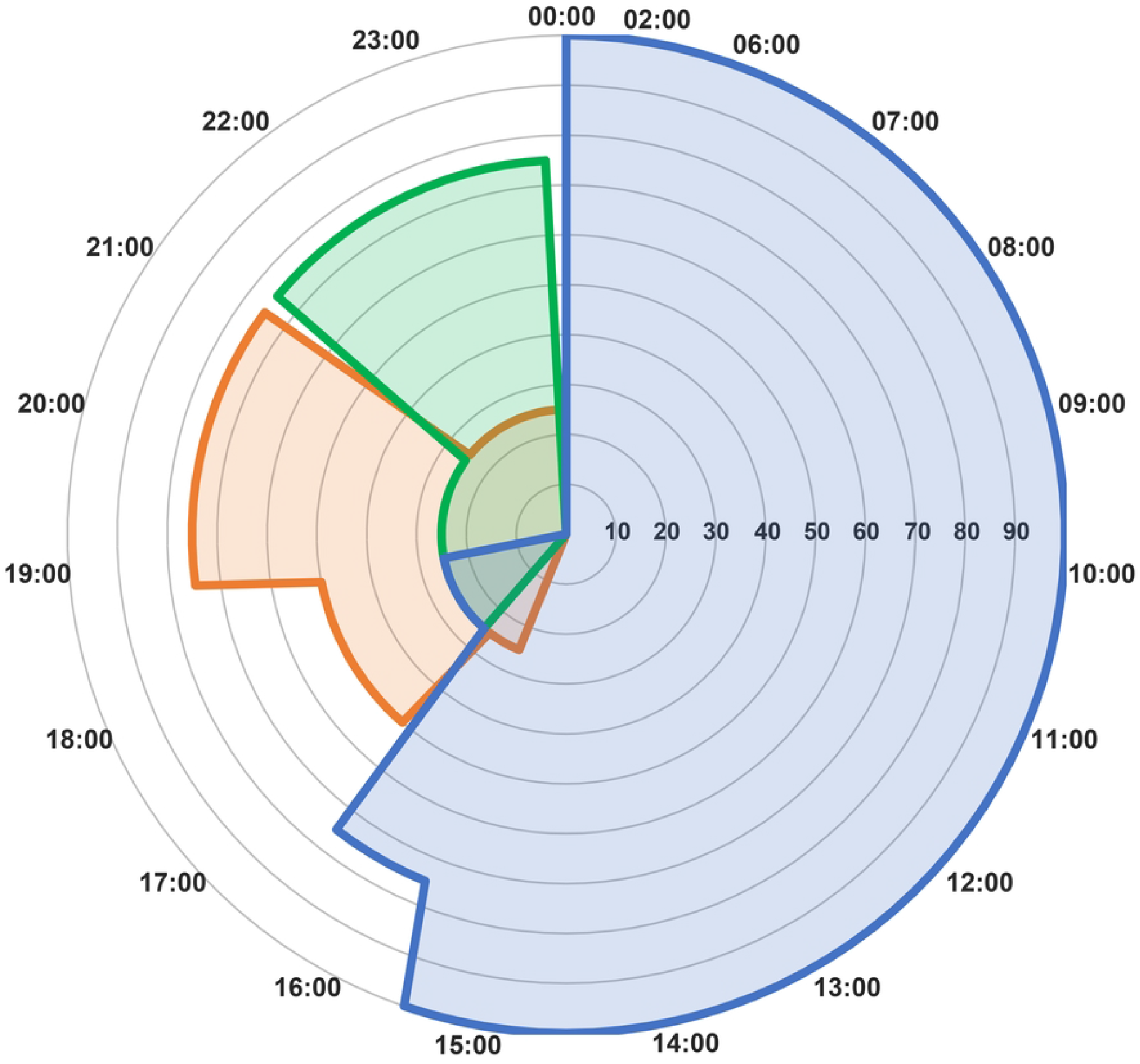
Floral phenology of *Annona macroprophyllata* in a 24-hour period. The female phase is activated during the afternoon, the species reaches its peak of receptivity during the night between 7 p.m. and 9 p.m. Then between nine and eleven at night, the flowers are mainly in the male phase. It is important to note that at each hour during anthesis, female and male flowers are synchronous in different proportions.

Throughout flower anthesis, low diversity and abundance of floral visitors were observed (20 species of arthropods). Four guilds were distinguished: pollinators, florivores, insect predators and random or non-specific visitors. Pollinators were insects that entered the pollination chamber during the female phase and remained inside the flower throughout anthesis (Fig. 3). 15 species of pollinators of the genera *Carpophilus* (Nitulidae) and *Colopterus* (Nitulidae) were found (67 individuals in total), of which *Carpophilus hemipterus* (20 individuals) was the most frequent, followed by *Carpophilus dimidiatus* (14 individuals), *Carpophilus lugubris* (5 individuals) and *Colopterus sp* (4 individuals). The highest number of pollinators was collected between 1600 and 2200 hours, which coincides with the highest peak of stigmatic receptivity among tres (Fig. 2). Among the other interactions observed, the only florivorous species detected was *Protographium epidaus* (Lepidoptera: Papilionidae), an insect specializing in feeding on *Annona* flowers. This species was observed in all its larval stages, consuming both young and mature petals. Also, two spiders (*Misumenoides* sp and *Hamataliwa* sp) were observed as insect predators and some fruit flies (*Drosophila*) and homopterans (*Membracis* sp.) were observed as non-specific visitors.

**Fig. 3.**
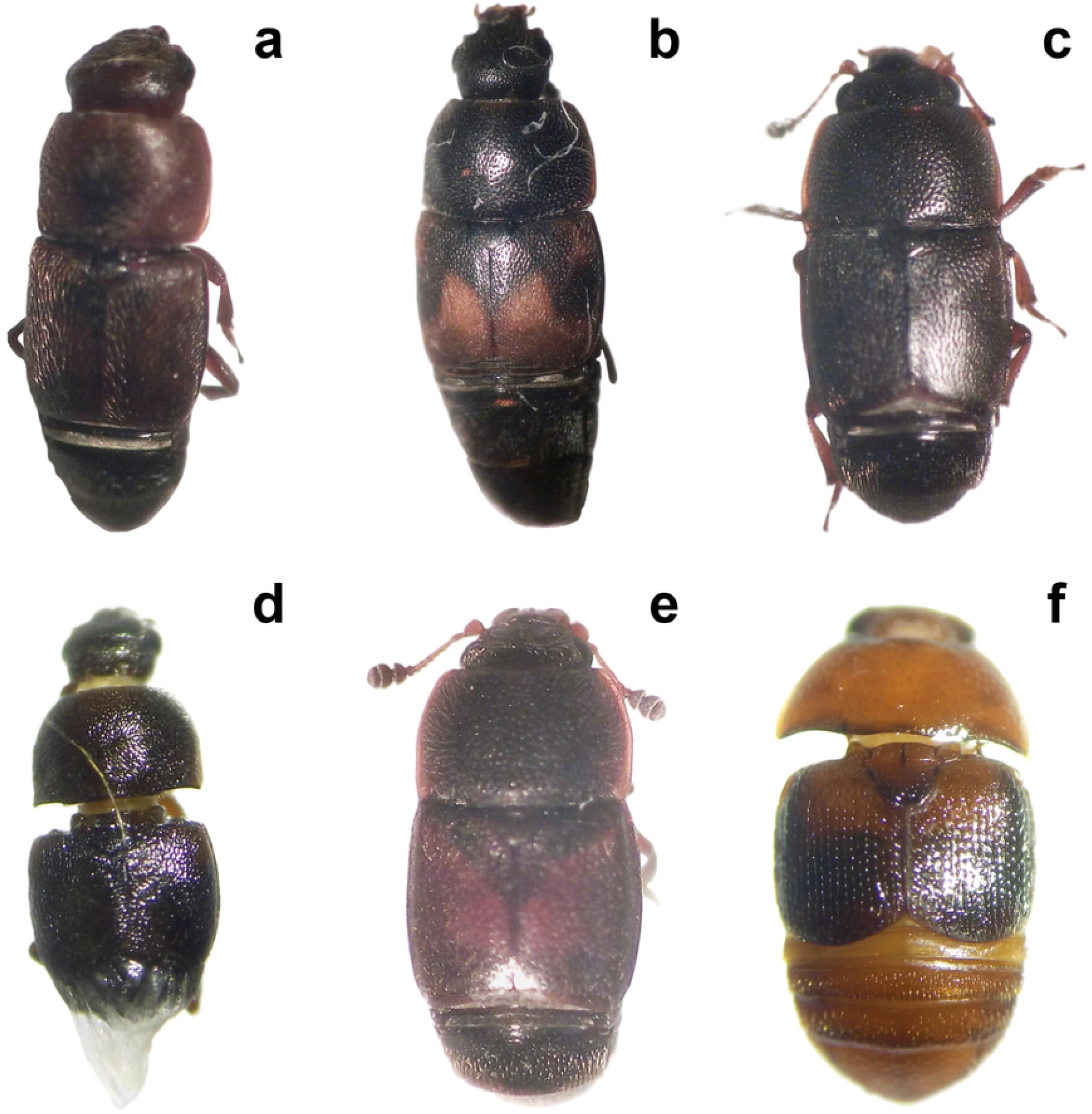
Nitulid pollinators of *Annona macroprophyllata*. Only the pollinators determined at the species level are illustrated.

### Flowers scents characterization

The comparison of floral scent diversity among flower phases showed that the sweet, fruity scents of female flowers are determined by the combination of esters, branched (or ramified) esters and carboxylic acids (Table 1). Thus, the scent of the female flowers could be indeed sweet, fruity and spicy garlicky. On the other hand, the fermented scents of the male flowers can be determined by the combination of furan derivatives, aldehydes, and cycloalkanes, which together can result in fermented, sour, green, or nutty odors (Table 1).

**Table 1.** Chemical compounds found in flowers in different reproductive phases (female and male). The compounds found have been linked to the scent of Annonaceae flowers.

| Retention time | Name of the compound | Molecule formula | Stigmatic flower (Female) | Staminate flower (Male) |
| --- | --- | --- | --- | --- |
| 1.398 | 4-Methyl-3-sulfanylpentanoic acid | C6H12O2S | x |  |
| 1.447 | 3-Methylpentyl propanoate | C9H18O2 | x |  |
| 2.348 | 3-Methyl-2-(2-oxopropyl)furan | C8H10O2 |  | x |
| 2.547 | Hexanal 5,5 dimethyl | C8H16O |  | x |
| 2.743 | 6-Hydroxy-2,2,6-trimethyl-3-(3-methylbut-2-enyl)cyclohexyl methyl acetate | C17H30O3 | x |  |
| 2.946 | 1,3-Dimethyl-5-isobutylcyclohexane | C12H24 |  | x |

## Discussion

### *Annona*-type floral syndrome

The floral phenology of *Annona macroprophyllata* is similar to that observed in other species of the genus [9]. Like most *Annona* species, *A. macroprophyllata* has receptive flowers at night (between 1900 and 2100 hours, with the peak of receptivity). Additionally, individuals of *A. macroprophyllata* can have both male and female flowers simultaneously, where geitonogamy is probably reduced by the fact that only a small proportion (25%) of the flowers are male during the peak receptivity in this species [1]. Something unusual about *A. macroprophyllata* is its extremely short anthesis (lasting only eight nighttime hours) and the likely overlap of the female and male phases in the same flower (the carpels do not fall off during pollen release). The short anthesis (one night) has also been reported for *Annona crassiflora* Mart., which is likely related to the circadian rhythms of its pollinators [8]. In the case of *A. macroprophyllata*, pollinators visit the flowers in the late afternoon and evening, at the beginning of the female phase. They remain inside the flower for about two hours and are gradually released during anthesis. After that, they visit another female flower, which indicates that they have a bimodal circadian rhythm [6]. The overlap of phases within the same flower has also been observed in *Annona exsucca* Dunal, with flowers appearing receptive even during pollen release [16]. However, in *A. macroprophyllata*, the carpels during the male phase are not covered by exudate. The exudate has been observed to facilitate pollen germination in Annonaceae [17], and its absence during male phase may prevent self-pollination in *A. macroprophyllata*.

In *Annona macroprophyllata*, pollination is carried out by small beetle insects of the Nitulidae family (Fig. 3). *Annona* species pollinated by small Nitulidae usually have small pollination chambers, nocturnal anthesis, and emit a sweet smell during stigmatic receptivity [8]. The floral characteristics of *A. macroprophyllata* match this pattern. Related species such as *A. cherimolla* and *A. squamosa* are also pollinated by small Nitulidae and exhibit similar floral traits to *A. macroprophyllata*, supporting the theory of a floral syndrome within the genus [1].

### Phenological changes in the chemical composition of flower odors in *Annona*

Pollinators of *Annona macroprophyllata* visit the flowers during the early female phase, which is signaled by the release of a sweet aroma and the formation of a pollination chamber. These insects remain inside the flower until they are released during the male phase. This behavior supports the hypothesis that pollinating insects are attracted only to female flowers. Upon their release during the male phase, these pollen-laden insects will likely visit only female flowers, thereby ensuring effective pollination [12]. Empirical observations of *Annona* species indicate that the flower’s scent varies between reproductive phases. However, no studies have characterised the scent composition during both phases in *Annona* flowers [11].

Our results demonstrate that the scent composition differs between the phases of *A. macroprophyllata* flowers. The scent of the female flower mimics the sweet smell of fruit, which usually attracts Nitulid beetles [5]. In contrast, the fermented odors of the male phase seem to repel them, contributing to their release or ensuring that they visit only female flowers. In other species of Annonaceae such as *Meiogyne virgata, Duguetia lanceolata, D. marcgraviana, D. neglecta*, and *D. pycnastera*, a scent shift from sweet to fermented during anthesis has also been documented [11]. The attraction-repulsion system in *Annona macroprophyllata* and other members of the Annonaceae family may represent an innovation independently acquired in each lineage. However, relatively few studies characterise the chemical compounds involved in floral odor throughout antesis [11]. Therefore, another possibility is that this system is a common floral attribute within the Annonaceae family.

## Acknowledgements

We would like to express our sincere gratitude to Guadalupe Amancio for her assistance in preparing the figures. Also thanks to Alejandro Zaldivar Riverón and Garet Powell for their help with the taxonomic identification of floral visitors. It’s important to note that this study is part of the first author’s Master’s program at the Universidad de Ciencias y Artes de Chiapas.

## Data availability

Data for this manuscript is available upon request through the corresponding authors.

